# Stability evolution as a major mechanism of human protein adaptation in response to viruses

**DOI:** 10.1101/2022.12.01.518739

**Authors:** Chenlu Di, Jesus Murga-Moreno, David Enard

**Author notes:** **Corresponding author:** David Enard.

## Abstract

Pathogens were a major driver of genetic adaptation during human evolution. Viruses in particular were a dominant driver of adaptation in the thousands of proteins that physically interact with viruses (VIPs for Virus-Interacting Proteins). This however poses a conundrum. The best understood cases of virus-driven adaptation in specialized immune antiviral factors or in host viral receptors are numerically vastly insufficient to explain abundant adaptations in VIPs. What adaptive mechanisms can then at least partly close this gap? VIPs tend to be broadly conserved proteins with conserved host native molecular functions. Because many amino acid changes in a protein can alter its stability –the balance between the folded and unfolded forms of a protein– without destroying conserved native activities, here we ask if stability evolution was an important mechanism of virus-driven human protein adaptation. Using predictions of protein stability changes based on Alphafold 2 structures and validated by multiple lines of evidence, we find that amino acid changes that altered stability experienced highly elevated adaptative evolution in VIPs, compared to changes with a weaker impact on stability. We further find that RNA viruses, rather DNA viruses, predominantly drove strong adaptation through stability changes in VIPs. We also observe that stability in immune antiviral VIPs preferentially evolved under directional selection. Conversely, stability in proviral VIPs needed by viruses evolved under compensatory evolution following viral epidemics. Together, these results suggest that stability evolution, and thus functional host protein abundance evolution, was a prominent mechanism of host protein adaptation during viral epidemics.

## Introduction

Virus-driven adaptation includes some of the most compelling and best understood examples of protein adaptation on a mechanistic level in human and other mammals. Host proteins with well understood adaptation in response to viruses notably include prominent immune antiviral proteins such as TRIM5 (Sawyer et al., 2005), PKR (Elde et al., 2009) and APOBEC3G (Compton et al., 2012; Sawyer et al., 2004; Yang et al., 2020) that directly attack viral molecules and trigger downstream molecular processes meant to destroy them. Other well understood cases include adaptive amino acid changes at the host-virus contact interface that decrease the binding affinity of viruses for their host cell surface receptors. The filovirus receptor NPC1 in bats is such an example, with one specific amino acid position in the contact interface explaining most of bat species-specific susceptibility (Ng et al., 2015). Interestingly however, viral receptors can also arbor many positively selected amino acid changes well outside of the contact interface with no identified mechanism to date to explain their adaptive nature (Enard et al., 2016; Uricchio et al., 2019). This is the case of ANPEP, a coronavirus receptor, where many selected sites well outside of the contact interface make it one of the most positively selected proteins in mammals (Enard *et al*., 2016).

More generally, recent evidence shows that, far from being restricted to specialized immune antiviral factors or viral receptors, adaptation has been abundant among thousands of currently known VIPs, including proviral ones, with a more than two-fold increase in adaptation compared to proteins that do no interact with viruses (noted non-VIPs) (Enard *et al*., 2016; Uricchio *et al*., 2019). VIPs in particular experienced highly increased rates of strong adaptation, where the viral selective pressure was likely intense and led to rapid fixation of the advantageous amino acid changes (Enard and Petrov, 2020; Uricchio *et al*., 2019). In VIPs, the surface contact interfaces with viruses only represent a very small proportion of all the amino acids, typically a few percent (Wierbowski et al., 2021). For example, the contact interface of ACE2 with SARS-CoV-2 represents only 21 amino acids, or 2.6% of ACE2 (Yan et al., 2020).

What protein evolution mechanisms then may explain widespread positive selection in VIPs not restricted to contact interfaces? What protein attributes changed that provided a selective advantage in response to viruses, while preserving the host native functions of VIPs? We can sort protein mutations into four non-exclusive categories that may be selected during viral epidemics: (i) mutations that affect protein conformation, (ii) mutations that alter the host-virus contact interface, (iii) mutations that change the host native molecular functions, and (iv) mutations that change the stability, i.e the abundance of the folded, functional form of VIPs (Dasmeh et al., 2013). Even though we do not exclude that any of these different mechanisms occurred in VIPs, it is important to consider that VIPs tend to be broadly conserved across mammals and beyond (Castellano, 2019). This is true irrespective of whether VIPs were discovered through low-throughput hypothesis-driven virology studies (75% with orthologs across mammals; Methods), or through high throughput mass-spectrometry assays that are blind to previous knowledge (74% with orthologs) (Batra et al., 2018; Jager et al., 2011; Shah et al., 2018; Watanabe et al., 2014). This excludes any difference in research attention and corresponding publications artificially biasing VIPs towards more conserved genes. This greater conservation might make repeated protein conformation changes less likely, because recurrent, notable conformational changes may be incompatible with the broad conservation of VIPs and their host native molecular functions (Konate et al., 2019; Lee et al., 2007; Radivojac et al., 2013). The latter might also limit adaptation through amino acid changes that modify the host native molecular activities of VIPs. Mutations at catalytic residues are very likely to disrupt these activities (Firnberg et al., 2014). In addition, similarly to contact interfaces, catalytic residues responsible for these activities only represent a small percentage of a protein (less than 5%), mainly restricted to protein surfaces (Nelson et al., 2013).

Conversely, protein stability changes may represent a good candidate mechanism for virus-driven adaptation. Protein stability can be defined as the thermodynamic balance between the folded functional form of a protein, and the unfolded, non-functional form that is typically targeted for degradation (Clausen et al., 2019). Protein stability changes are quantified as ΔΔG, defined as the change in the thermodynamic quantity ΔG, the Gibbs free energy (in kcal/mol) that determines the stability of a protein. Amino acid changes with positive ΔΔG are destabilizing, negative ones are stabilizing. A change of ΔΔG of one kcal/mol roughly corresponds to a fivefold change in folded protein abundance at body temperature (Dasmeh *et al*., 2013; Serohijos et al., 2013). Protein stability is a major determinant of the abundance of the functional form of a protein in cells, and many known disease-causing variants destabilize and decrease the abundance of essential proteins (Nielsen et al., 2017; Scheller et al., 2019; Stein et al., 2019). We can then imagine how VIP stability changes may be advantageous during a viral epidemic. For example, lower stability and thus lower abundance of a proviral factor required by a virus to replicate may be advantageous for the host.

Experimental data, notably from the disease variants literature (Stein *et al*., 2019) and from protein design studies (Goldenzweig and Fleishman, 2018), shows that many amino acid changes in many parts of a protein can change stability (Li et al., 2020; Serohijos and Shakhnovich, 2014). The large number of possible stability-altering amino acid changes may thus in theory match the large number of adaptive amino acid changes observed in VIPs. Furthermore, stability evolution may be easier in otherwise conserved proteins (Dasmeh *et al*., 2013), including VIPs; a large pool of possible stability-altering amino acid changes might (i) happen outside of evolutionarily conserved active sites of VIPs responsible for conserved host native functions, and (ii) a large pool might make compensatory evolution easier in the case where a host native function is no longer optimal after a change in folded protein abundance (see below, Compensatory stability evolution in proviral VIPs). Finally, stability changes are particularly likely for amino-acid changes that occur in the buried parts of protein structures, and below we observe that many adaptive amino acid changes in VIPs occurred within the buried part of VIPs (Results below).

Here, we focus on protein stability changes as a possible mechanism of adaptation in response to viral epidemics. We find that amino acid substitutions in human evolution that altered protein stability substantially were much more likely to be adaptive in VIPs, compared to amino acid changes that changed stability to a lesser extent. We further find that VIPs have experienced more adaptation in the buried than in the outer parts of their protein structure. This is in agreement with the fact that changes of buried residues are more likely to affect protein stability, and we confirm indeed that the elevated adaptation in the buried part of VIPs is driven by those amino acid changes that modify protein stability. We further observe that antiviral and proviral VIPs have experienced stability-driven adaptation, but in different ways. Immune antiviral VIPs whose main function is to impede viruses have overall experienced directional stability evolution, likely as a result of changing optima depending on shifting pathogen pressures over evolutionary time. Conversely, most proviral VIPs do not have immune functions but have many broadly conserved non-immune host native functions under stabilizing selection. As expected, likely due to these conserved native functions, non-immune proviral VIPs experienced predominantly compensatory stability evolution. Together, our result suggest that protein stability evolution may have been an important mechanism of host adaptation in response to viruses. Our results further suggest a model where further compensatory evolution may occur after viral epidemics, to bring proviral VIPs back to their initial, and perhaps more optimal stability in the absence of viral selective pressure.

## Results

To test if protein stability was a determinant of adaptation in VIPs, we use a recent version of the McDonald-Kreitman test (McDonald and Kreitman, 1991) called ABC-MK (Uricchio *et al*., 2019) to estimate the proportion of amino acid substitutions that were adaptive among those that significantly altered stability, compared to those that did not during human evolution since divergence with chimpanzees. McDonald-Kreitman approaches estimate the percentage of nonsynonymous substitutions that were adaptive by contrasting the total observed number of nonsynonymous substitutions with what this number would be under neutrality, if adaptation had not occurred. This neutral expectation can be derived from the Site-Frequency-Spectrum of present non-synonymous variants, while at the same time controlling for past fluctuations of the mutation rate by contrasting nonsynonymous and synonymous substitutions and variants from the same coding sequences (Uricchio *et al*., 2019). We use ABC-MK with coding sequence substitutions that occurred specifically in the human branch since divergence with chimpanzees (Methods), and variants from the 1,000 Genome projects groups located in Africa (Genomes Project et al., 2015), as described in (Uricchio *et al*., 2019). Notably, ABC-MK has the ability to distinguish between weak and strong adaptation (Uricchio *et al*., 2019) (Methods), and here we use this functionality as it was previously shown that viruses drive particularly strong adaptation (Enard and Petrov, 2020; Uricchio *et al*., 2019).

We estimate stability changes caused by amino acid substitutions and variants (Figure 1A,B) with the computational method Thermonet (Li *et al*., 2020) used on high confidence Alphafold 2 protein structures (Jumper et al., 2021) (available at https://alphafold.ebi.ac.uk/; Methods). Thermonet uses a deep convolutional network trained on experimental stability data to predict the stability changes caused by amino acid substitutions or variants within a given protein structure. Experimental measures of stability changes are not currently available beyond a limited number of human proteins (Pancotti et al., 2022).

**Figure 1.**
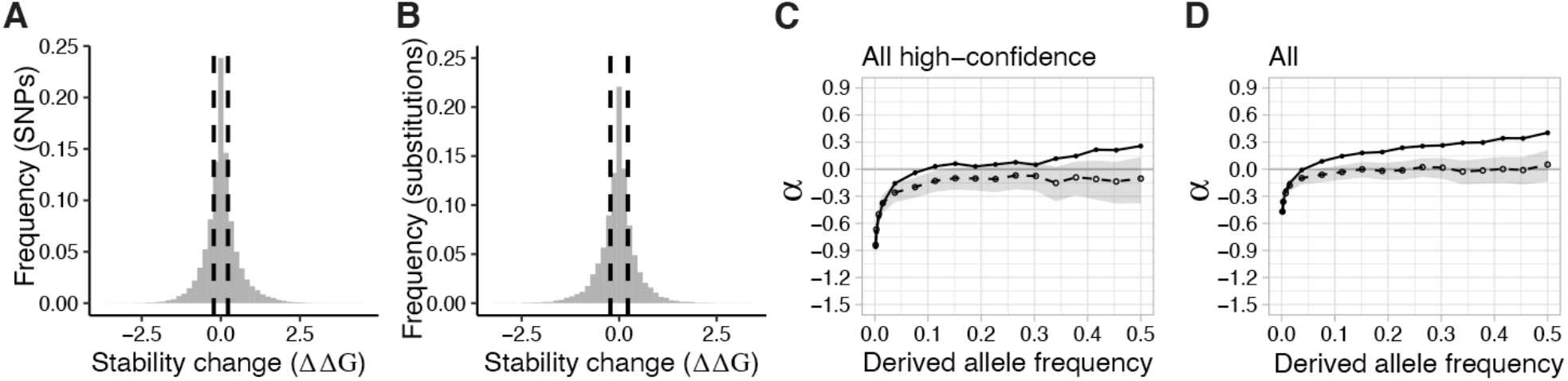
Distributions of stability changes and adaptation in VIPs and control non-VIPs. A) Distribution of Thermonet estimated ΔΔG values for human nonsynonymous variants (Methods). The dashed lines represent the ΔΔG=-0.225 and ΔΔG=0.225 limits. B) Distribution of Thermonet estimated ΔΔG values for human nonsynonymous fixed substitutions (Methods). C) α-curves used by ABC-MK to estimate the proportion a of adaptive nonsynonymous substitutions (y-axis; Methods) using only variants and substitutions from high confidence Alphafold 2 residues (pLDDT≥70) with stability predictions (for the nonsynonymous ones; Methods) in VIPs (continuous curve) and control non-VIPs (dashed line for the average and grey area for the 95% confidence interval). D) Same as C) but using all variants and substitutions known at VIPs and control non-VIPs, not just the ones restricted at high confidence Alphafold 2 residues.

Although Thermonet provides computational estimates, it was recently shown to have good, balanced performance when benchmarked with new experimental stability data that was not used for its convolutional network training (Pancotti *et al*., 2022). We further validate Thermonet estimates in multiple ways. First, Thermonet correctly identifies a known destabilizing genetic variant (R105G) in antiviral VIP APOBEC3H as strongly destabilizing (stability change ΔΔG = 0.71 kcal/mol) (Chesarino and Emerman, 2020). Second and most importantly, we find strong, highly significant evidence of compensatory evolution of protein stability in non-immune proviral VIPs with multiple amino acid changes, where it is the most expected (see below). It would not be possible to observe such compensatory evolution if Thermonet estimates were not sufficiently correlated with the actual stability changes that occurred during human protein evolution.

Out of ~5,500 VIPs known to date (Table S1), 2,900 have high confidence Alphafold 2 structures and can be compared with 5,700 non-VIPs also with high confidence structures (Table S1; Methods). These include only VIPs and non-VIPs with orthologs across mammals (Methods) since we previously showed that viruses increase adaptation more specifically in VIPs that are conserved across mammals and beyond (Castellano, 2019). This also limits the risk of confounding due to gene age (Moutinho et al., 2022). We compare VIPs with non-VIPs to highlight the evolutionary patterns that are specific to VIPs (Figure 1C,D).

Here, we specifically ask if stability-altering substitutions have (i) experienced more positive selection in VIPs than in non-VIPs, and (ii) more positive selection than substitutions that altered stability to a lesser extent. We do not compare VIPs with any non-VIPs, but match VIPs with control non-VIPs that look like VIPs in ways other than interacting with viruses, that could affect adaptation and confound the comparison (Methods). The matching is done with a previously described bootstrap procedure (Di et al., 2021; Enard and Petrov, 2020) and includes many potential confounders (Methods). We then estimate how significant increased adaptation is in VIPs by repeating the measurements of adaptation in sets of the same size as the VIPs set, but made of randomly sampled VIPs and non-VIPs. This effectively estimates an unbiased false discovery rate by generating null distributions of estimates of adaptation expected if there was no impact of interactions with viruses (FDR; Methods).

Considering only substitutions and variants with predicted stability changes (Methods), the 2,900 VIPs with good Alphafold 2 structures have experienced 3.6 times more adaptation than control non-VIPs, with 27.5% of all amino acid substitutions estimated to be adaptive vs. only 7.4% for control non-VIPs (Figure 1C; the proportion of adaptive substitutions is noted a). This includes 19% of strongly selected substitutions in VIPs, compared to only 3% in control non-VIPs. Considering all substitutions and variants including those with no stability change prediction (Methods), the 2,900 VIPs have experienced 3.5 times more adaptation than control non-VIPs, with 35% of all amino acid substitutions estimated to be adaptive vs. only 10% for control non-VIPs (Figure 1D). This includes 29% of strongly selected substitutions in VIPs compared to only 4% in control non-VIPs.

### Abundant stability-changing, adaptive substitutions in VIPs

In order to study the impact of protein stability on adaptation, we split nonsynonymous variants and fixed substitutions in the human branch according to stability changes (given by their |ΔΔG| absolute value in kcal/mol) into two groups: those below the variants’ absolute stability change median (|ΔΔG|<0.225), and those above (|ΔΔG|>0.225). This also happens to roughly correspond to a transition point in the leptokurtic distribution of ΔΔG among variants or substitutions (Figure 1A,B), between a large concentration of SNPs around ΔΔG=0 and more spread out, symmetric tails on both sides. We do not try to distinguish between destabilizing (ΔΔG>0) and stabilizing (ΔΔG<0) variants that decrease or increase protein stability, respectively, but instead we use the absolute |ΔΔG|. Indeed, VIPs can be proviral or antiviral, making it tempting to assign an expectation of what stability change direction should be adaptive. One might expect that it is advantageous to increase the stability and therefore increase the abundance of an antiviral VIP indefinitely. This may however have adverse effects, notably in the case of antiviral VIPs related to inflammation (Yong et al., 2022). More importantly, we found through a large, multi-year effort of manual curation of 4,477 virology publications that antiviral VIPs for a virus are very frequently subverted and made proviral by the same or other viruses (Table S1, Methods), thus making it difficult to assign a specific expected direction of adaptive ΔΔG. Retasking by viruses to accomplish proviral steps was also recently noticed to occur even with very prominent antiviral factors (King and Mehle, 2022; Tran et al., 2020). Finally, assigning an adaptive ΔΔG direction might also be made difficult by the fact that there might be compensatory evolution of stability, especially in proviral VIPs that have important, conserved non-immune functions in the host (immune antiviral VIPs are more specialized in attacking viruses and likely less impeded by other native host functions, see below).

Thus, we estimate the rate of adaptive substitutions with ABC-MK for just two categories, all substitutions with absolute stability changes below, and all substitutions above the variants’ absolute median for |ΔΔG|, respectively (Figure 1A,B). We cut the data this way to compare two groups of equal sizes, and thus similarly powered comparisons between VIPs and non-VIPs. We call the group above the median the Large Stability Changes group (LSC), and the group below the Small Stability Changes group (SSC). We only include variants and substitutions at amino acids with high Alphafold 2 structural prediction confidence (Alphafold 2 accuracy score pLDDT≥70; Methods).

We find much higher rates of adaptation for the LSC group than for the SSC group. ABC-MK estimates that 34% or 250 of the 737 LSC substitutions were advantageous in VIPs, versus only 15% or 125 of the 832 SSC substitutions (Figure 2A,B). Thus, a substantial majority (67% or 250/375) of advantageous nonsynonymous substitutions with ΔΔG predictions in VIPs altered protein stability. The estimated 34% of adaptive LSCs in VIPs is also much higher than the estimated 10% in sets built randomly sampling both VIPs and control non-VIPs to estimate a false discovery rate (FDR=3*10^−4^; Methods). In contrast, VIPs only have a marginally higher adaptive percentage of SSCs compared to random sets (15% vs. 7% respectively, FDR=0.04). By comparing adaptive proportions in VIPs and control non-VIPs, we can also estimate the amount of adaptation that can be attributed to interactions with viruses. In control non-VIP sets, 8% LSC substitutions were advantageous on average, more than four times less than in VIPs. A total of 34% minus 8% (26%), or 192 of the 737 LSC substitutions in VIPs, can thus be attributed to virus-driven adaptation. SSC substitutions were also 8% advantageous in control non-VIPs, implying 58 (15%-8%) of the 832 SSC substitutions in VIPs can be attributed to viruses. Thus, 77% (192/(192+58)) of the adaptative substitutions attributable to viruses in VIPs changed stability above the |ΔΔG| median.

**Figure 2.**
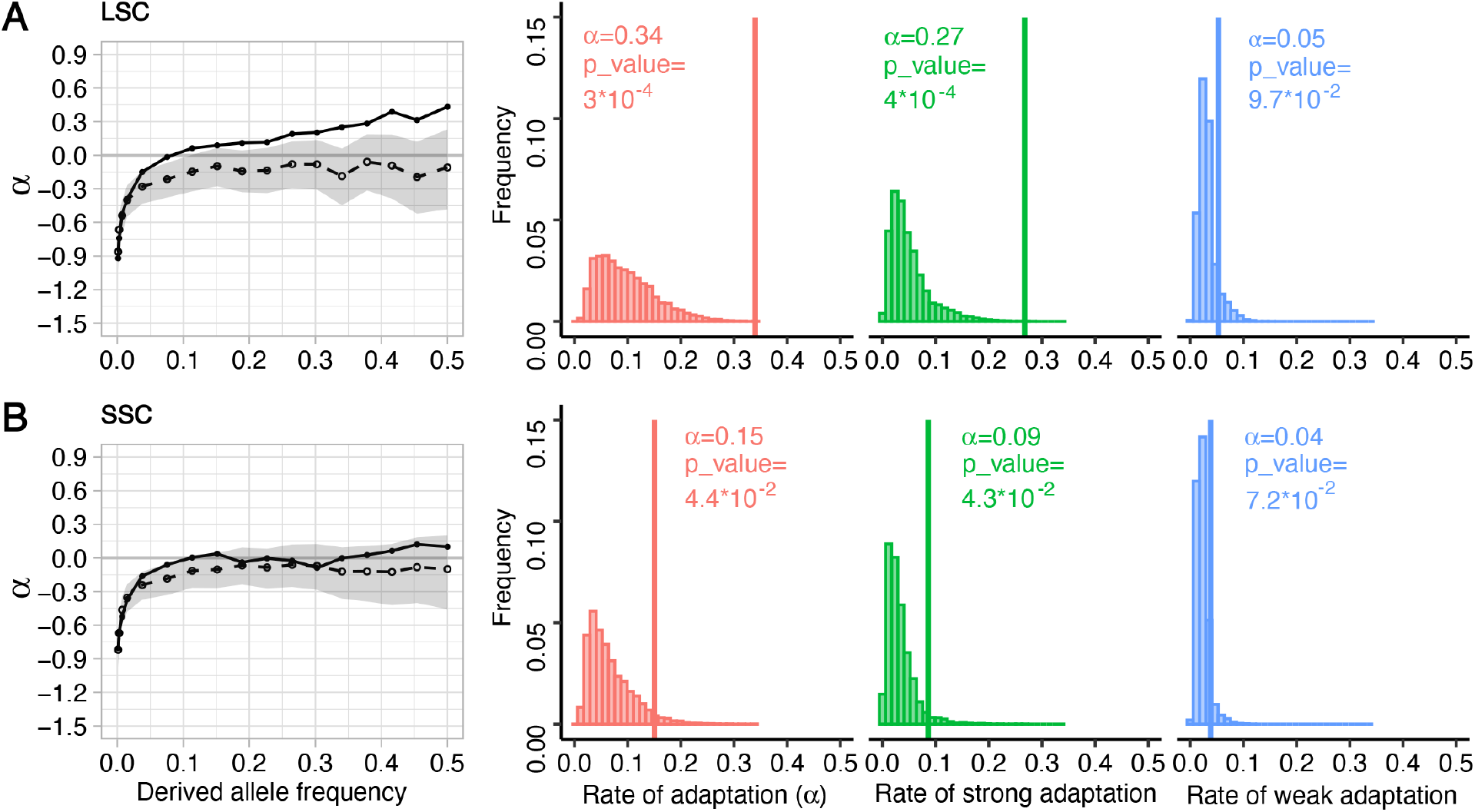
Stronger adaptation through large stability changes in VIPs. We compared adaptation among LSC substitutions (A) with adaptation among SSC substitutions (B). The α-curves used by ABC-MK to estimate α are as in Figure 1C. The red graph shows the total α in VIPs (vertical line) vs. the expected null distribution when randomly sampling VIPs and control non-VIPs. Green graph: same as red but only for strong adaptation as measured by ABC-MK. Blue graph: same as red but only for weak adaptation as measured by ABC-MK.

ABC-MK also has the ability to distinguish between strong and weak past protein adaptation (Methods). We find that in VIPs, it is in particular the rate of strong adaptation that is increased among LSC substitutions, with an estimated 27% of substitutions being strongly advantageous, vs. only 5% in random sets and 3% in control non-VIPs (Figure 2A; FDR=4*10^−4^). In VIPs SSC substitutions are 9% strongly advantageous compared to 4% in control non-VIPs. Using the same logic as above, this means that 177 LSC and 42 SSC substitutions can be attributed to strong, virus-driven adaptation, respectively, with 81% of strong virus-driven adaptation then involving stability changes above the |ΔΔG| median. This corresponds to the expected evolutionary pattern if adaptative evolution with LSCs in VIPs was indeed driven by viruses, since we previously showed that virus-driven adaptation was disproportionately strong adaptation (Enard and Petrov, 2020; Uricchio *et al*., 2019).

### Stability, rather than other protein attributes explains increased VIP adaptation at buried residues

We further assess potential confounding protein attributes that might be the primary selected attributes, instead of stability. Specifically, the effect of a given residue on stability is known to depend strongly on the position of the residue in the protein structure. More hydrophobic residues closer to the buried core of proteins are known to contribute disproportionately to protein stability (Nick Pace et al., 2014). In agreement with this, LSCs tend to be more buried, further from the structural surface of VIPs than SSCs. Using Relative Solvent Accessibility (RSA) measured by DSSP (Kabsch and Sander, 1983) (Methods) as a measure of how buried or exposed to the surface a given protein residue is, we observe that LSCs have an overall median RSA of 0.23, vs. 0.4 for SSCs (see also Figure S1). This however raises the possibility that buried residues in VIPs have elevated adaptation regardless of their impact on stability. VIPs might experience more adaptation at buried residues because changes in protein conformation and/or changes in allostery, where functional signals are transmitted through protein structure (Nelson *et al*., 2013; Swain and Gierasch, 2006), were the primary adaptive protein attributes. The increased rate of adaptation of LSCs might then only be a secondary, bystander effect of the fact that substitutions in buried parts of proteins tend to also affect protein stability to a greater extent. Accordingly, we find that VIPs experienced more adaptive substitutions below than above the overall median RSA (RSA=0.32) in their structures, with 39% below or 262 adaptive substitutions, vs. 19% above or 173 adaptive substitutions estimated by ABC-MK, respectively (Figure 3A,B). The increased adaptation below the median RSA strongly sets VIPs apart from control non-VIPs (random sets FDR=2*10^−4^; Figure 3B), and this is even more the case for strong adaptation (Figure 3B). Notably, LSCs were less adaptive in high RSA (Figures 3C and S2A) than in small RSA (Figures 3D and S2B) parts of VIPs, which excludes that adaptive LSCs overall can represent a bystander effect of adaptation at contact interfaces or molecular activities located at the protein surface.

**Figure 3.**
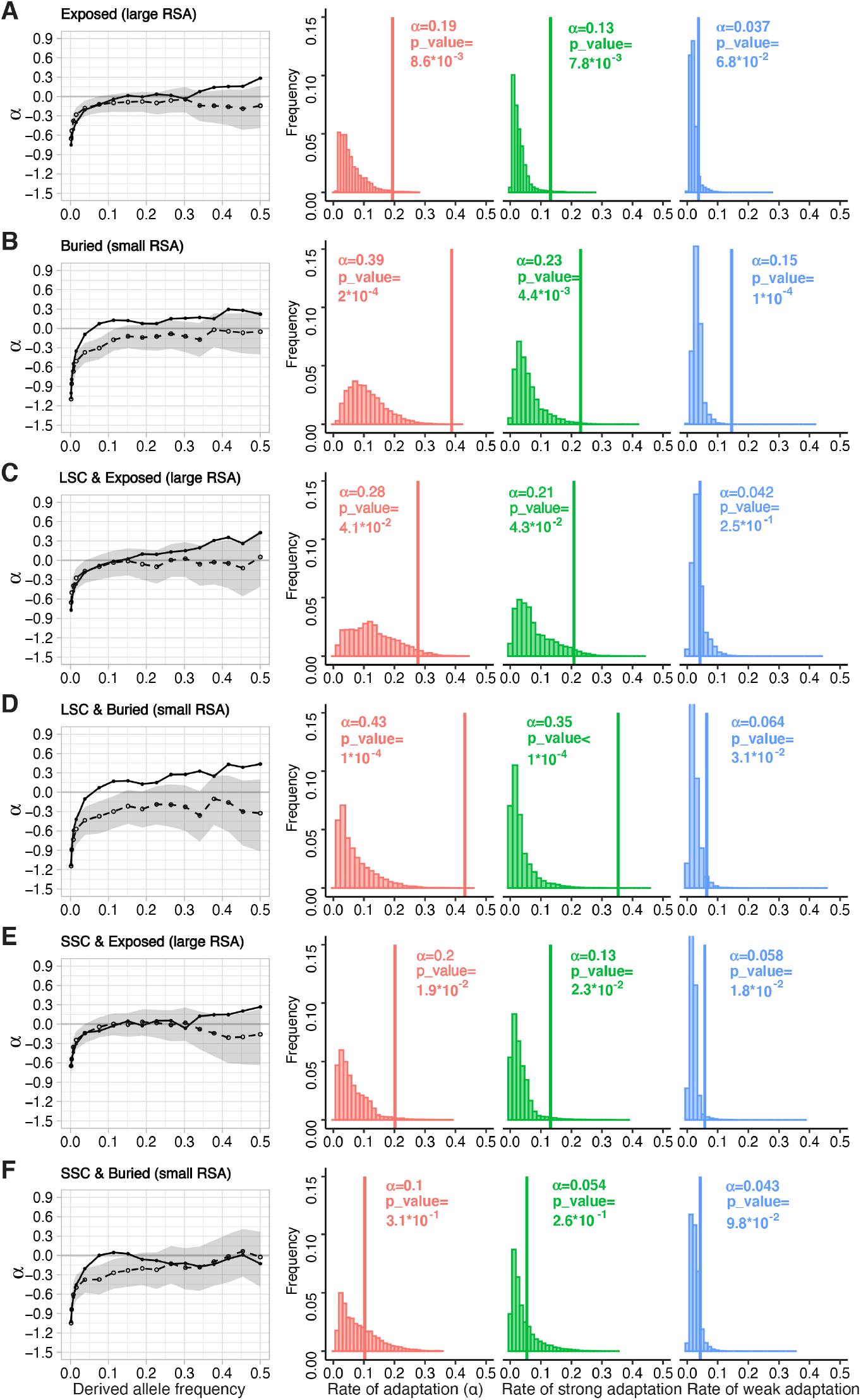
Adaptation as a function of position in the protein structure. Legend same as Figure 2. A) adaptation (LSC+SSC) in more exposed parts of VIPs and control non-VIPs with RSA≥0.32, the median for LSC and SSC combined. B) same as A) but for more buried parts with RSA<0.32. C) LSC adaptation above the LSCs’ RSA median of 0.23. D) LSC adaptation below the LSCs’ RSA median of 0.23. E) SSC adaptation above the SSCs’ RSA median of 0.4. F) SSC adaptation below the SSCs’ RSA median of 0.4. All four subgroups represented in C, D, E and F have separating RSA medians such that the four subgroups have similar sizes (and thus variance and power to test hypotheses).

We further observe that the increased adaptation in the low RSA parts of VIPs is strongly dependent on protein stability. While LSCs below their RSA median or the SSCs’ RSA median (0.23 and 0.4, respectively) have strongly elevated adaptation compared to random sets (Figures 3D and S2), all SSCs in VIPs below a 0.4 RSA only have 10% adaptive substitutions, a percentage that is not different from random FDR sets (FDR=0.31) and lower than the 20% adaptive SSCs in high RSA parts of VIPs (Figure 3E,F). Together, these results suggest that protein stability changes are the primary driver of increased adaptation, especially strong adaptation, in more buried, lower RSA parts of VIPs, rather than changes of protein conformation and/or allostery. This further narrows down the possible mechanistic explanations to protein stability, since more buried parts of VIPs are also less likely to include contact interfaces or active catalytic pockets typically found at the surface of proteins.

### Increased adaptation through large stability changes in more RNA than DNA viruses

Having found broad patterns across VIPs, we then estimate which viruses have a particularly increased associated percentage of adaptive LSC substitutions. We previously found that RNA viruses whose genomes are coded by RNA, rather than DNA viruses, have driven particularly strong and abundant selection at their respective VIPs during human evolution (Enard and Petrov, 2018; 2020; Souilmi et al., 2021). If abundant adaptive evolution of LSCs in VIPs is indeed a hallmark of virus-driven adaptation, we should then be able to observe that this is particularly the case for VIPs of RNA viruses. In agreement with this prediction, we find seven RNA viruses out of nine tested with significantly increased adaptive LSCs in their specific VIPs (compared to random FDR sets, Table 1), compared to only one of the six tested DNA viruses, Kaposi’s sarcoma Herpesvirus KSHV (Table 1). Five of the seven significant RNA viruses have VIPs with very strongly elevated percentages of adaptive LSCs at or above 50%, including coronaviruses (72%, Figure 4A,B), rhinovirus RV-B14 (84%, Figure 4C,D), Dengue Virus (58%, Figure S3A), Human Immunodeficiency Virus HIV (54%, Figure S3B) and Influenza A Virus (50%, Figure S3C). For all significant viruses except HIV, the strong elevation of adaptive LSCs mostly reflects strong adaptation (Table 1, Figures 4 and S3). In the VIPs of coronaviruses, dengue virus, Hepatitis C virus, HIV, rhinovirus RV-B14 and KSHV, the difference in adaptation between LSCs and SSCs is extreme (Table 1 and Figures 4 and S3). All VIPs that interact only with RNA viruses had a significant elevation of adaptive LSCs (40%, Table 1), while all VIPs that interact only with DNA viruses did not (10%, Table 1). Together, these results show that RNA viruses were the predominant drivers of strong adaptation through large stability changes in VIPs.

**Table 1.** LSC and SSC adaptation for the VIPs and control non-VIPs of 15 different viruses. α is the total alpha. α_s_: strong adaptation. α_w_: weak adaptation. DN: number of nonsynonymous substitutions (LSC left or SSC right) in the VIPs of each virus. DN-α: corresponding estimated number of adaptive nonsynonymous substitutions. Upper rows: nine RNA viruses. Middle rows: six DNA viruses. Lower rows: VIPs that interact only with RNA viruses and VIPs that interact only with DNA viruses. Light orange: random shuffling of VIPs and control non-VIPs FDR<0.05. Dark orange: FDR<0.001.

|  | LSCs |  |  |  |  | SSCs |  |  |  |  |
| --- | --- | --- | --- | --- | --- | --- | --- | --- | --- | --- |
| | $\alpha$ | $\alpha_s$ | $\alpha_w$ | DN | DN-α | $\alpha$ | $\alpha_s$ | $\alpha_w$ | DN | DN-α |
| coronaviruses | 0.72 | 0.58 | 0.09 | 118 | 84.43 | 0.05 | 0.02 | 0.02 | 105 | 5.35 |
| DENV | 0.58 | 0.46 | 0.07 | 135 | 77.93 | 0.08 | 0.03 | 0.02 | 150 | 11.39 |
| EBOV | 0.18 | 0.05 | 0.08 | 41 | 7.29 | 0.53 | 0.31 | 0.14 | 48 | 25.40 |
| HCV | 0.45 | 0.21 | 0.18 | 86 | 38.30 | 0.05 | 0.02 | 0.01 | 111 | 5.19 |
| HIV | 0.54 | 0.23 | 0.27 | 158 | 84.82 | 0.05 | 0.02 | 0.02 | 194 | 10.35 |
| IAV | 0.50 | 0.38 | 0.07 | 157 | 78.87 | 0.47 | 0.34 | 0.09 | 192 | 90.39 |
| RHINOV | 0.84 | 0.76 | 0.07 | 25 | 21.02 | 0.04 | 0.01 | 0.01 | 26 | 0.91 |
| WNV | 0.18 | 0.16 | 0.13 | 44 | 12.65 | 0.36 | 0.19 | 0.05 | 56 | 20.00 |
| ZIKV | 0.45 | 0.24 | 0.16 | 93 | 41.71 | 0.49 | 0.39 | 0.08 | 119 | 58.81 |
| EBV | 0.18 | 0.09 | 0.03 | 139 | 24.96 | 0.18 | 0.08 | 0.06 | 131 | 24.05 |
| HBV | 0.06 | 0.05 | 0.15 | 57 | 8.35 | 0.06 | 0.02 | 0.01 | 76 | 4.17 |
| HPV | 0.20 | 0.13 | 0.03 | 141 | 28.33 | 0.39 | 0.28 | 0.07 | 138 | 54.45 |
| HSV | 0.04 | 0.02 | 0.01 | 46 | 1.94 | 0.61 | 0.45 | 0.09 | 64 | 38.79 |
| KSHV | 0.55 | 0.37 | 0.13 | 108 | 59.46 | 0.04 | 0.02 | 0.01 | 89 | 3.86 |
| VACV | 0.44 | 0.27 | 0.05 | 47 | 20.68 | 0.03 | 0.01 | 0.01 | 63 | 1.93 |
| RNA only | 0.40 | 0.31 | 0.06 | 286 | 114.16 | 0.04 | 0.02 | 0.01 | 339 | 13.73 |
| DNA only | 0.11 | 0.04 | 0.04 | 137 | 14.45 | 0.04 | 0.01 | 0.02 | 139 | 5.02 |

**Figure 4.**
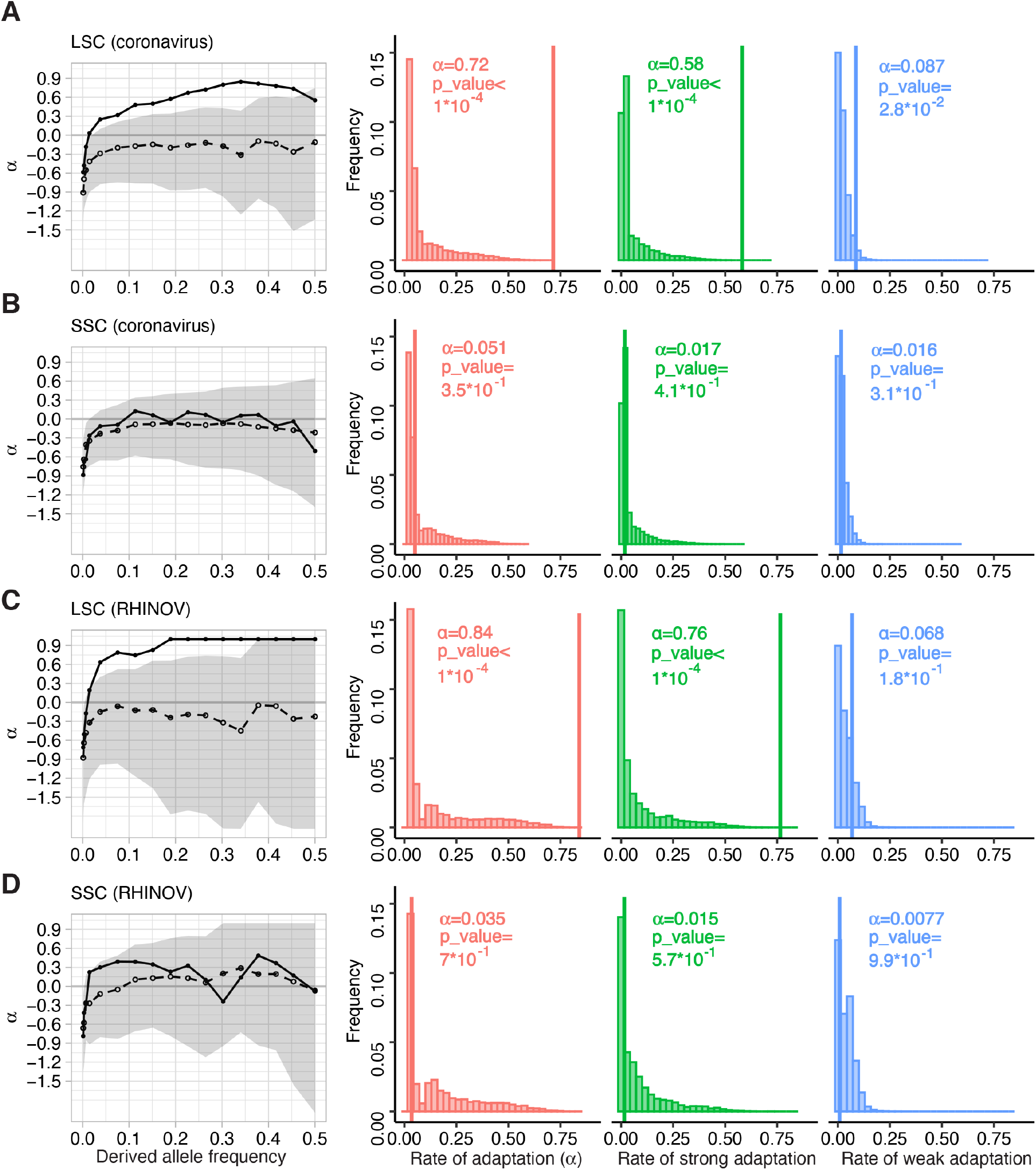
Adaptation in large or small stability changes for the VIPs of two RNA viruses with high LSC adaptation. Legend same as Figure 2. A) ABC-MK results for 448 coronavirus VIPs and control non-VIPs for LSCs. B) Same but for SSCs. C) ABC-MK results for 164 rhinovirus RV-B14 VIPs for LSCs. D) Same but for SSCs. The numbers of VIPs for each virus type correspond to the number of VIPs with enough high confidence Alphafold 2 residues (Methods) and with orthologs across mammals (Methods).

### Directional evolution in immune antiviral VIPs, and compensatory evolution of stability in non-immune proviral VIPs

Because we use computational estimations from Thermonet, we seek to further validate these estimations by testing predictions about the expected evolution of protein stability of different functional classes of VIPs, that can only be verified if Thermonet’s estimations are of sufficient quality. Different VIPs have very diverse native functions in the host (Enard and Petrov, 2018; 2020), but can still be sorted in a number of categories depending on their functional effects on viruses. In particular, many VIPs can be classified as immune antiviral VIPs that actively engage viral molecules to directly degrade, or target them for degradation, or activate degradation pathways (Table S1). On the other end of the functional spectrum, many VIPs do not have any immune function and are in fact proviral factors that assist viruses when the latter hijack their molecular functions to complete multiple steps of their replication cycle (Table S1). For example, viruses subvert host transcription regulators to activate the expression of their own viral genes during infection (Chau et al., 2008; Scoggin et al., 2001; Shen et al., 2022). We predict that immune antiviral and non-immune proviral VIPs should have experienced different patterns of protein stability evolution. In addition to uncovering new evolutionary patterns of host adaptation to viruses, verifying these predictions would also provide strong additional validation of the quality of the ΔΔG estimated by Thermonet.

We predict that immune antiviral VIPs should have experienced predominantly directional stability evolution, while non-immune proviral VIPs should have experienced predominantly compensatory evolution of stability. Indeed, the main, often highly specialized function of immune antiviral VIPs is to engage with viruses, with limited pleiotropic constraints in their way to often evolve very rapidly (Elde *et al*., 2009; Sawyer *et al*., 2004; Sawyer *et al*., 2005). Their incessant arms race with viruses suggests that stability as an adaptive protein attribute, might have then evolved under directional selection, with stability optima changing over time together with the ever-changing landscape of pathogenic viruses. Conversely, non-immune proviral VIPs have non-immune, conserved native functions in the host. Thus, in any given non-immune proviral VIP with stability itself under stabilizing selection, adaptation during viral epidemics might take away protein stability from its usual optimum to complete native host functions, when there is no viral selective pressure to temporarily shift that optimum. This predicts that compensatory evolution of protein stability should then occur in non-immune proviral VIPs after viral epidemics, in the form of amino acid changes that tend to bring stability back closer to its pre-epidemic level (Chaurasia and Dutheil, 2022). This prediction has the advantage that it can be tested in a straightforward way, provided that Thermonet estimations are of sufficient quality. If compensatory stability evolution occurred in a non-immune proviral VIP, then we expect that the sum of stability changes caused by each amino acid change in this VIP should be closer to zero than expected by chance (regardless of the chronological order of these changes, which is unknown). Stability changes in this VIP should indeed compensate each other, i.e. should tend to cancel each other out. Conversely, we might observe the opposite for an immune antiviral VIP under directional stability evolution, with the sum of stability changes caused by each amino acid substitution being further from zero than expected by chance, if the direction of selection was preferentially toward increasing, or toward decreasing stability.

This is straightforward to look at because we can compare the sum of stability changes associated with each substitution in a VIP with the null expected distribution under neither stabilizing compensatory nor directional selection. We get the null expected distribution by randomly shuffling estimates of protein stability changes between genes included in the analysis (Methods), while also preserving the absolute average stability change per substitution in each VIP with shuffled stability changes (Methods). Crucially, the comparison with null expectations also provides a test of the quality of Thermonet ΔΔG predictions; we do not expect any departure from random expectations if the predictions are too far from the actual effects of the substitutions on protein stability.

The difficulty is then to identify immune antiviral, and non-immune proviral VIPs. To know which VIPs fall into these two categories, we conducted an extensive manual curation of 4,477 virology articles on VIPs, looking for the reported functional effects of VIPs on the replication cycle of a wide range of viruses (Methods). Through this effort we were able to identify 772 immune antiviral VIPs and 875 non-immune proviral VIPs as of October 2022 (Table S1; Methods).

We must further consider a few important potential limitations. In the millions of years since divergence with chimpanzees (a short amount of time for protein divergence), most VIPs with amino acid substitutions have accumulated only one such substitution and are therefore not appropriate for testing directional or compensatory evolution that imply multiple substitutions. Thus, we restrict this analysis to 80 immune antiviral VIPs and 52 non-immune proviral VIPs with two or more amino acid substitutions in the human branch (Table S1). We further consider that it might take more than one additional substitution to complete an episode of compensatory evolution after an epidemic, or more than two overall substitutions to start seeing a clear unidirectional pattern of directional selection. Thus, in addition to the test with VIPs with two or more nonsynonymous substitutions, we also further restrict our analysis to VIPs with three or more substitutions, in particular with the expectation that the signatures of compensatory evolution should be more visible in this subset of VIPs. We also restrict the test to VIPs with a minimal number of increasingly large absolute stability changes. Indeed, VIPs with no pronounced stability change may be entirely intolerant to such changes, or may not require compensatory evolution at all in the first place.

Using this specific, increasingly restrictive testing design, we first find that the 80 immune antiviral VIPs with at least two substitutions (Table S1) have predominantly experienced directional stability evolution. On average across these VIPs, the sum of stability changes is 1.17 fold further from zero than expected by chance (stability shuffling test P=2.5*10^−3^; Methods). As predicted, this departure tends to become more pronounced when focusing on immune antiviral VIPs with a larger number of increasing high stability changes (Figure 5A,B). Individual antiviral VIPs with a strong signature of directional stability evolution notably include the APOBEC3G antiviral factor, albeit with marginal statistical significance (Table S1; sum of stability changes two times higher than expected, stability shuffling test P=0.07). Although immune antiviral VIPs have an overall collective trend toward directional selection, we notice a few notable exceptions. TRIM5 has a signature of compensatory, not directional stability evolution (Table S1). The sum of the six stability changes in TRIM5 is 10.7 times closer to zero than expected by chance (stability shuffling test P=0.049). Similarly, the prominent antiviral factor OAS1 has three stability changes with a signature of compensatory evolution (sum 20.1 times closer to zero than expected by chance, stability shuffling test P=0.028).

**Figure 5.**
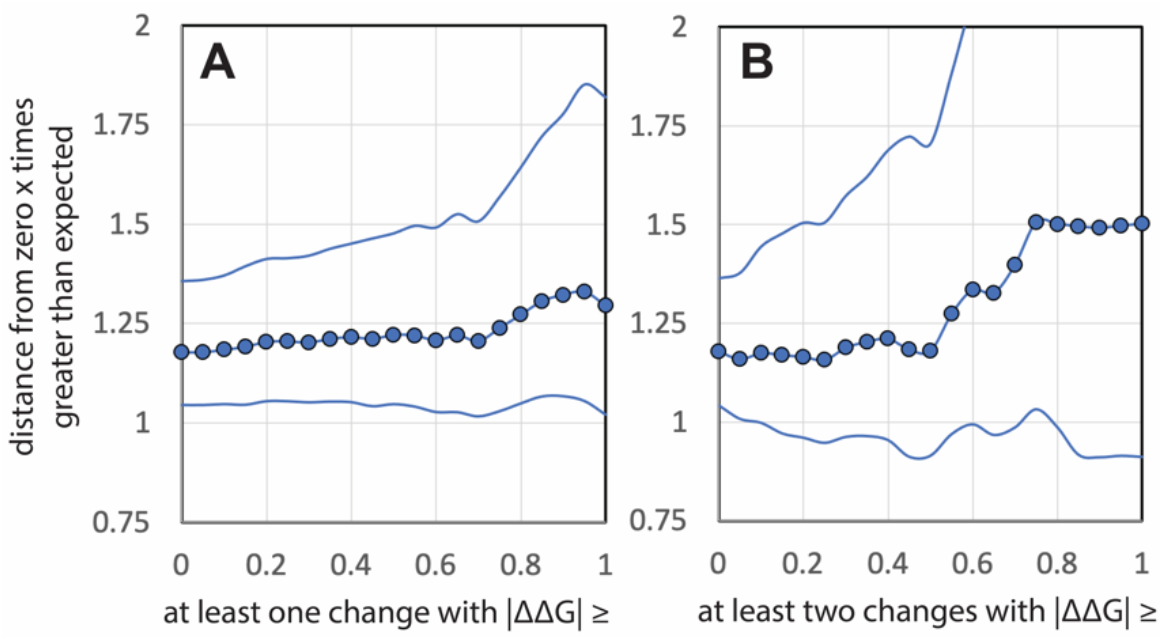
Directional evolution in immune antiviral VIPs. The y-axis represents how many times further from zero the average of the per-VIP |sum| of stability changes is compared to random expectations. Central curve: compared to average random expectations over 1,000,000 random iterations (Methods). Lower curve: 95% confidence interval lower boundary. Upper curve: 95% confidence interval upper boundary. The x-axis represents the |ΔΔG| threshold to restrict the shuffling test only to groups of immune antiviral VIPs each with (A) at least two substitutions in total including at least one at or above the |ΔΔG| threshold, or (B) at least two substitutions in total including at least two at or above the |ΔΔG| threshold.

But compensatory stability evolution is most visible in non-immune proviral VIPs. For these VIPs we observe an overall trend of the sum of stability changes being on average closer to zero than expected by chance (Figure 6, y-axis). Overall, the sum is 1.2 times closer to zero than expected by chance in the 52 non-immune proviral VIPs with at least two nonsynonymous substitutions (stability shuffling test P=0.02; Methods). This trend becomes much more pronounced when restricting the test to non-immune proviral VIPs with three or more nonsynomymous substitutions, and when restricting the test to VIPs with an increasing number of more pronounced stability changes (Figure 6A,B,C). For example, the sum of stability changes is on average 2.5 times closer to zero than expected in the 16 non-immune proviral VIPs with at least three nonsynonymous substitutions including two with |ΔΔG|≥0.2 (Figure 6C and Table S1, stability shuffling test P=1.6*10^−5^). This strong increase of the compensatory evolution signature compared to when including any nonsynonymous substitutions makes sense, given that many non-immune VIPs with weaker |ΔΔG| substitutions likely do not need compensatory evolution in the first place.

**Figure 6.**
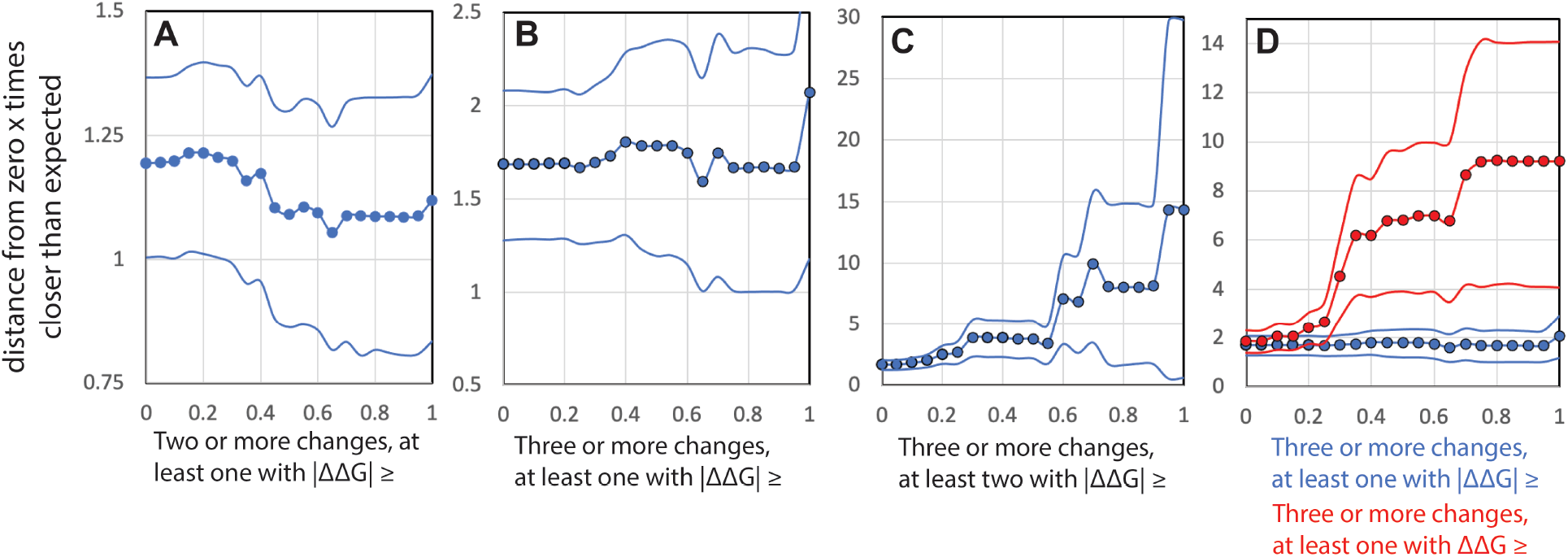
Compensatory evolution in non-immune proviral VIPs. The y-axis represents how many times closer to zero the average of the per-VIP |sum| of stability changes is compared to random expectations. Central curve: compared to average random expectations over 1,000,000 random iterations (Methods). Lower curve: 95% confidence interval lower boundary. Upper curve: 95% confidence interval upper boundary. The x-axis represents the |ΔΔG| threshold to restrict the shuffling test only to groups of non-immune proviral VIPs each with (A) at least two substitutions in total including at least one at or above the |ΔΔG| threshold, or (B) at least three substitutions in total including at least one at or above the |ΔΔG| threshold, or (C) at least three substitutions in total including at least two at or above the |ΔΔG| threshold. The blue curves in (D) are the same as in (B). The red curves in (D) represent how much closer the average of the per-VIP |sum| of stability changes is compared to random expectations in VIPs with at least three substitutions in total, including at least one at or above the destabilizing ΔΔG threshold on the x-axis. Note the very different ranges of the y-axis for A,B,C and D.

Having clarified the possible impact of compensatory evolution in non-immune proviral VIPs, we further test one last prediction. Because proviral VIPs benefit the viruses that subvert them, strongly destabilizing substitutions that strongly decrease the abundance of non-immune proviral VIPs should be particularly advantageous during viral epidemics, and may also require similarly strong compensatory evolution afterwards. Thus, we expect that compensatory evolution should be particularly visible in non-immune proviral VIPs with at least one strongly destabilizing substitution, compared to non-immune proviral VIPs with at least one substitution affecting stability similarly but in either direction, stabilizing or destabilizing. When we compare these two situations, we find a much stronger signal of compensatory evolution in non-immune proviral VIPs specifically with at least one strongly destabilizing substitution (Figure 6D). This signal increases when restricting the test to VIPs with at least one increasingly destabilizing substitution (Figure 6D). This supports the predicted model where strongly destabilizing substitutions made proviral VIPs less available to viruses during epidemics, followed by compensatory evolution with stabilizing substitutions.

Together, these results highlight important differences in the stability evolution of two functional types of VIPs. The predominant signature of directional selection in immune antiviral VIPs, and most importantly the strong signature of compensatory evolution in non-immune proviral VIPs, cannot be expected if the Thermonet predictions were not of good, sufficient quality. Indeed, it is difficult to see how the sum of stability change predictions would be so much closer to zero than expected in non-immune proviral VIPs, if those predictions were far from the actual, real effect of the corresponding amino acid changes on protein stability. It would also be hard to explain how the compensatory evolution signature could be so much stronger in non-immune proviral VIPs with strongly destabilizing substitutions. This further provides strong support for the Thermonet ΔΔG predictions.

## Discussion

Using Thermonet predictions of protein stability changes, we found that such changes were likely an important mechanism of virus-driven host adaptation. Although the computational nature of these predictions may raise doubt about their quality, it is important to reiterate that poor predictions not much better than random would not have allowed us to observe a large adaptation difference between LSCs and SSCs in VIPs, or the marked differences between immune antiviral and non-immune proviral VIPs (directional vs. compensatory evolution). Specifically, our results with non-immune proviral VIPs suggests an evolutionary model with positive selection during and after viral epidemics, with changes in stability during epidemics that are later compensated back to pre-epidemic stability levels that are likely more optimal for native host functions. This evolutionary model however requires further investigation. In particular, we do not have access to the chronological order of substitutions that would allow to further test it.

Together with previous studies, our results further highlight a common theme of host adaptation through changes in RNA and/or protein abundance of genes that interact functionally with pathogens. We previously found that strong selection during an ancient coronavirus or related virus epidemic in East Asia predominantly occurred at or in close linkage to eQTLs of coronavirus VIPs (Souilmi *et al*., 2021). Similarly, Klunk et al. recently found that the most strongly selected variants during the Black Death are associated with changes in the expression level of immune genes (Klunk et al., 2022). There is also evidence that adaptive introgression in response to viruses, from Neanderthals to Eurasian modern human ancestors, involved Neanderthal variants that affect gene expression (Enard and Petrov, 2018; Nedelec et al., 2016; Quach et al., 2016).

Importantly however, our results do not exclude that mechanisms other than VIP stability changes also have played an important role in virus-driven protein host adaptation. Adaptive evolution in viral receptors might cut epidemics short by blocking viruses from infecting cells altogether. Similarly, adaptation in prominent antiviral factors such as TRIM5 or APOBECs may have had an oversized impact compared to VIPs not specialized in attacking viruses (Sawyer *et al*., 2004; Sawyer *et al*., 2005). Quantifying the relative quantitative contributions of different mechanisms of host adaptation is however complicated by the fact that the location of the vast majority of physical contact interfaces between viruses and VIPs are currently unknown. Further understanding of virus-driven adaptation will likely require a better knowledge of these interfaces. That said, we still do not expect that adaptation at contact interfaces may be able to fully explain the abundant adaptation we observe for large stability changes, in a scenario where these would only be a secondary bystander effect. Indeed, we found strong stability-dependent adaptation in more buried parts of the structures of VIPs. This however does not exclude the possibility of adaptive changes in binding between viruses and VIPs also due to allosteric, distance effects of buried adaptive amino-acid substitutions, through conformational and/or structural flexibility changes. It is also important to note that we only focused on parts of proteins that are well-structured with high Alphafold 2 confidence scores (Methods). Intrinsically disordered protein segments are known to result in poor structure confidence scores, and were thus completely excluded from our analysis. Whether intrinsically disordered proteins or protein segments also participate substantially to virus-driven adaptation remains an open question (Lou et al., 2016).

Our results suggest that in the future, identifying the biological mechanisms involved in virus-driven adaptation might enable more discriminant detection of ancient viral epidemics, where the involvement of a specific virus in past epidemics may be recognized not only through an overall increase in adaptation in its specific VIPs, but more specifically through an increase in adaptative evolution of specific mechanistic attributes such as protein stability. In that respect, we find that RNA viruses clearly stand out during human evolution compared to DNA viruses. Taken together, our results show that in addition to the detailed functional study of specific gene candidates by evolutionary virologists, the study of quantitative patterns of adaptive evolution in VIPs as a whole group can provide new insights on the functional evolutionary changes that gave hosts a fighting chance against repeated viral epidemics.

## Methods

### Identifying human protein coding genes with orthologs across mammals

We previously found that viruses increase adaptation specifically in VIPs with orthologs across mammals (Castellano, 2019). We therefore restrict all analyses to VIPs and non-VIPs with orthologs across mammals. We use an updated list of Ensembl v99 human genes (Cunningham et al., 2019) with orthologs found in at least 251 out of 261 mammalian genome assemblies. These 261 assemblies were extracted from NCBI Genome (https://www.ncbi.nlm.nih.gov/genome/) and are the deposited assemblies that had a N50 contig size of at least 30kb as of July 2021, in order to limit the number of truncated genes. Ensembl human protein coding genes are selected as mammals-wide orthologs if they have best-reciprocal hits using the largest number of identical nucleotide hits from Blat alignments (Kent, 2002), with at least 251 out of the 261 genome assemblies (to account for the fact that orthologs in some species may have not been sequenced, and located in assembly gaps). This process finds 13,495 such human Ensembl v99 genes with orthologs across mammals (Table S1). The list of mammalian species and the corresponding assembly versions are provided in Table S1.

### Thermonet predictions with Alphafold structures

We use the Thermonet software to predict the ΔΔG caused by specific amino acid changes in VIPs and non-VIPs. Thermonet uses a convolutional neural network to make ΔΔG predictions. Thermonet’s neural network is trained using images of the biophysical properties of the close three-dimensional environment of the amino-acid change location. The neural network was trained first on experimental datasets of ΔΔG measurements. Because Thermonet uses the three-dimensional local environment, it requires a protein structure as input. To run Thermonet we chose to use public Alphafold v2 structures (from https://alphafold.ebi.ac.uk/) rather than experimental structures because using only available human experimental structures would have strongly limited the number of VIPs and the statistical power of the analysis. Alphafold 2 however generates structures very close to the experimental ones when the latter are available to use by Alphafold as input (Jumper *et al*., 2021). This means that for those proteins with experimental structures, we expect very little difference when using the Alphafold structures.

The advantage of Alphafold is then also that it provides good quality predictions for a substantially larger number of human proteins than are experimentally available (Jumper *et al*., 2021). This is because Alphafold accurately predicts structures made of local folds that are well represented in its input database (Jumper *et al*., 2021). Nevertheless, Alphafold still fails to properly predict a subset of proteins or parts of proteins that then have a mixture of well and poorly predicted local structure regions. Note that this is particularly true for proteins or parts of proteins that are intrinsically disordered and thus do not have one single structure to predict in the first place.

Fortunately, Alphafold provides a site-by-site confidence score, noted pLDDT (Jumper *et al*., 2021). The pLDDT score has been shown by comparison with experimental structures to strongly predicts the per-residue structure accuracy. The pLDDT score varies from zero to 100, with 100 indicating the most accurate per-residue structural prediction possible. A pLDDT score above 70 is usually indicative of a highly accurate structure prediction. For this reason, in our analysis we only use Thermonet ΔΔG predictions at sites with a pLDDT equal to or greater than 70. We also only use Alphafold structures with 50% or more sites with pLDDT≥70. In total 76% of the VIPs and non-VIPs that we compare have such high quality Alphafold 2 structures (same % for VIPs and non-VIPs), for a total of 2,909 and 5707 non-VIPs. In addition, to avoid any confounding effect of discrepancies between the accuracy of Alphafold structures between VIPs and non-VIPs, we match VIPs with control non-VIPs with similar average per-residue pLDDT and percentage of sites with pLDDT≥70 (see below, VIPs and control non-VIPs with matching confounding factors).

Using the filters described above, we use 86,244 and 10,337 Thermonet ΔΔG predictions for coding variants and substitutions, respectively. Figure 1 represents the distribution of the predicted ΔΔG for variants (Figure 1A) and substitutions (Figure 1B). We run Thermonet using the amino acid change from the ancestral to the derived amino-acid, from the ancestral to the fixed human amino acid for substitutions, and from the ancestral to the derived allele for variants. Finally, it is also important to note that Alphafold only provides publicly the structures of the canonical coding sequence of each protein coding gene, but not for their other isoforms. Here we thus use only the corresponding Ensembl v100 canonical coding sequences with an Alphafold structure, which excludes variants and substitutions in other isoforms that do not overlap with the canonical isoform.

### Variants and substitutions data for ABC-MK

We use ABC-MK as previously extensively described in (Uricchio *et al*., 2019), using the human coding variants and substitutions dataset also described in the same publication. The main difference is that we split nonsynonymous variants and substitutions into two groups, noted LSCs and SSCs in the main text, according to their predicted ΔΔG that we then run ABC-MK on separately. We separate the two groups according to the variants absolute median of |ΔΔG| (0.225) in the 2,909 VIPs included in the analysis. We do this so that the groups have similar amounts of information, and thus the same variance of ABC-MK estimates of the proportion of selected substitutions. This proportion is usually noted α, calculated as α=1-(PN*DS)/(PS*DN), where PN and PS are the numbers of nonsynonymous and synonymous variants, respectively, and DN and DS are the numbers of nonsynonymous and synonymous fixed substitutions, respectively. The classic MK test simply uses this calculation of α. ABC-MK uses a more complex approach by computing the α-curve, which is the curve of a measured specifically for bins of derived allele frequencies, as for example in Figure 1C. This is required to account for segregating non-synonymous deleterious variants among other things (Uricchio *et al*., 2019). ABC-MK uses Approximate Bayesian Computation to match the observed α-curve with the best-fitting ones among many analytically predicted α-curves. Each analytical α-curve is generated for millions of combinations of varying distributions and amounts of deleterious, weakly advantageous, and strongly advantageous mutations. The difference between weakly and strongly advantageous variants that is exploited by ABC-MK is that weakly advantageous mutations do not go to fixation so fast that their contribution to nonsynonymous polymorphism is negligeable as it is for strongly advantageous variants (Uricchio *et al*., 2019). This affects the shape of the α-curve, especially at higher derived allele frequencies where weakly advantageous variants tend to segregate before eventually reaching fixation (selective sweeps tend to have a long pre-fixation phase after the faster exponential one). This translates into a downward trend of the α-curve at higher frequencies that can be detected by ABC-MK. It is important to note that another possible cause of a downward trend of the α-curve at higher frequencies is mispolarization, where low frequency derived nonsynonymous alleles may be mistaken for high frequency derived ones. This can happen when a nucleotide site experienced a substitution in the human branch, but subsequently experienced a mutation back to the initial ancestral nucleotide. This new derived allele will then be mistakenly annotated as the ancestral (Hernandez et al., 2007). Hernandez et al. have shown that in the human genome this issue affects derived alleles with a frequency greater than 0.7. We therefore run ABC-MK using nonsynonymous and synonymous variants with a derived allele frequency less than 0.7. It is also important to mention that ABC-MK uses the shape of the α-curve at low derived allele frequencies to estimate the distribution of deleterious fitness effects (Urrichio et al.). An important last detail about how we run ABC-MK is that because as described above, we only use amino acids with a pLDDT score at or above 70, we only use the synonymous variants and substitutions in the corresponding codons. We use the same set of synonymous variants and substitutions as a neutral reference when we run ABC-MK either with LSCs or SSCs, since focusing on smaller subsets of nonsynonymous changes (either LSCs or SSCs) is readily accounted for by the PN over DN ratio in the equation α=1-(*PN*\*DS)/(*DN*\*PS). ABC-MK is available at https://github.com/jmurga/MKtest.jl.

### Controlling for confounding factors when matching VIPs and non-VIPs

To highlight evolutionary patterns of adaptation that are specific to VIPs, we compare them with sets of control non-VIPs that have been matched with VIPs so that the former have the same average values of confounding factors as the latter. Here confounding factors are factors other than physical interaction with viruses that in principle might affect adaptation, and thus explain differences between VIPs and non-VIPs instead of physical interactions with viruses. We know for example that VIPs tend to be much more highly expressed at the mRNA level than non-VIPs. Higher expression might hypothetically be associated with increased adaptation, and thus explain the increased adaptation in VIPs rather than interactions with viruses. We match control non-VIPs with VIPs using a bootstrap procedure that was already extensively previously described (Di *et al*., 2021; Enard and Petrov, 2020). In total we match 17 potential confounding factors between VIPs and control non-VIPs:

-Ensembl canonical coding sequence length, since they correspond to the isoform used by Alphafold.
- the average GC content for each coding sequence.
- the average GC1 content at the first codon nucleotide position for each coding sequence.
- the average GC2 content at the second codon nucleotide position for each coding sequence.
- the average GC3 content at the third codon nucleotide position for each coding sequence. GC1, GC2, and GC3 control for possible differences in GC content between nonsynonymous and synonymous sites that might distort the α-curve.
- average GTEx v8 (Consortium, 2020) TPM (Transcripts Per Million) mRNA expression across 53 tissues (in log base 2).
- average GTEx v8 TPM mRNA expression in lymphocytes (in log base 2).
- average GTEx v8 TPM mRNA expression in testis (in log base 2).
- the number of protein-protein interactions (in log base 2) in the human protein interaction network (Luisi et al., 2015).
- the proportion of immune genes as annotated with the Gene Ontology terms GO:0002376 (immune system process), GO:0006952 (defense response) and/or GO:0006955 (immune response) as of May 2020 (Gene Ontology, 2015).
- the recombination rate (Halldorsson et al., 2019) in 500kb windows centered on genes, to account for potential mutational biases related to recombination such as biased gene conversion that could differentially affect synonymous and nonsynonymous sites with different GC content. We use large 500kb windows because they better represent the long-term recombination rate in a given genomic window compared to smaller windows.
- McVicker’s B value (McVicker et al., 2009), a measure of background selection that we used to account for the recent prevalence of segregating deleterious variants in the genomic environment surrounding a coding sequence and that could affect adaptation (Di *et al*., 2021).
- the density of GERP conserved elements (Davydov et al., 2010) in 50 kb and 500 kb windows centered on genes, to further account for the possible prevalence of segregating deleterious variants in the genomic environment surrounding a coding sequence.
- the proportion of amino acids in the Alphafold v2 structure with a pLDDT score of 50 or above.
- the proportion of amino acids in the Alphafold v2 structure with a pLDDT score of 70 or above.
- the average pLDDT score in the entire Alphafold structure.

### Estimation of Relative Solvent Accessibilty

Relative Solvent Accessibility (RSA) provides a measure of how exposed at the surface or buried close to the protein core specific amino acids are in the Alphafold structures. We measure RSA using DSSP (Kabsch and Sander, 1983).

### Functional annotation of VIPs

As part of an ongoing effort to annotate the functional impacts of VIPs on viruses, we have to date manually curated 4,477 virology articles from Pubmed to collect their proviral and/or antiviral effects (Table S1), as reported by the virology experiments described in those virology articles. We also annotated if the proviral and antiviral effects occurred through an immune function of the VIPs, either directly reported by the virology articles or because of the fact that the corresponding VIPs are annotated with the Gene Ontology terms GO:0002376 (immune system process), GO:0006952 (defense response) and/or GO:0006955 (immune response) as of May 2020. Through this annotation we identified 772 VIPs with an immune antiviral impact, and 875 VIPs with a non-immune, proviral impact (Table S1). Interestingly, we also found that 321 of the 772 antiviral immune VIPs for at least one virus, also have proviral effects for the same or different viruses. A detailed inspection of such cases shows that it happens for example when an expressed antiviral immune VIP has a molecular activity involved in the immune response that is subverted by viruses. For example, the antiviral VIP CREBBP is an important transcription activator involved in interferon beta production (Qu et al., 2021), that is exploited by the Human T-cell Leukemia virus to activate the transcription of its own viral genes (Scoggin *et al*., 2001). Antiviral immune VIPs expressed during infection are likely good targets for proviral repurposing by viruses, due to their broad availability at the precise time of need of their molecular activities by viruses.

### Directional and compensatory evolution of VIP stability

To detect directional or compensatory evolution of stability in VIPs, we design a random shuffling test based on the sum of stability changes in individual VIPs, then averaged over VIPs with multiple amino acid changes tested together. For example, in non-immune proviral VIPs we expect each individual VIP with multiple amino acid changes to have stability changes that tend to cancel out each other, thus resulting in a sum of stability changes closer to zero than expected by chance. As a statistic we use the average across VIPs of the absolute value of the sum of stability changes in each VIP. We then compare this average with its random expectation. This random expectation is generated as follows: for each VIP with a number x of stability changes, we randomly sample x stability changes from the entire pool of all predicted stability changes. We iterate this random sampling for each given VIP, until the randomly sampled stability changes have an average |ΔΔG| that matches the observed average |ΔΔG| for this VIP (plus or minus 2%). We do this to account for the fact that different VIPs may have different spreads of their distribution of possible stability changes, which could affect the null random expected sum of stability changes for each VIP. For the whole set of tested VIPs, we iterate this process 1,000,000 times to determine the average random expectation and the statistical significance of any observed departure of the real average sum of stability changes from this expectation. The sets of VIPs tested together must fulfill a number of pre-requisites, such as a minimal total number of stability changes, and a minimum number of those stability changes having their individual |ΔΔG| above a fixed threshold. For example, for non-immune proviral VIPs, we use these pre-requisites with increasingly stringent thresholds supposed to restrict the test to a number of VIPs where signatures of compensatory evolution are expected to be more visible. Indeed, compensatory evolution is more likely to have occurred in VIPs with a larger number of stability changes (compensatory changes had the chance to occur in the first place), and with a minimum number of large |ΔΔG| changes (compensatory evolution is likely required only when large |ΔΔG| changes occurred in the first place). We represent in Figure 5 the ratio of the observed average absolute value of the sum of stability changes, divided by the average random expectation (and the inverse for Figure 6) and its 95% confidence intervals upper and lower values generated by the 1,000,000 random samplings.

## Acknowledgements

We thank Mike Barker, Joanna Masel and Ryan Gutenkunst for their comments on the manuscript. David Enard is funded by NIH NIGMS MIRA R35GM142677.

