## Supplementary Figures for "Stability evolution as a major mechanism of human protein adaptation in response to viruses"

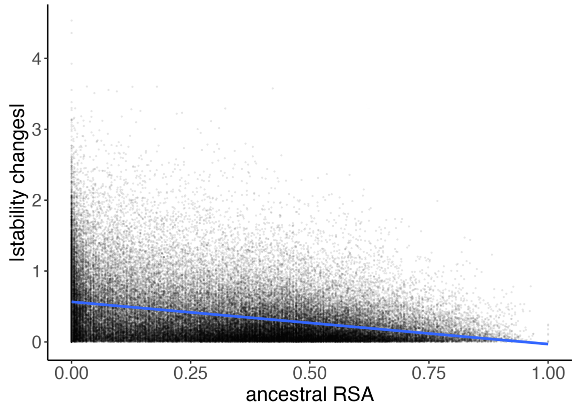


**Figure S1. Correlation between |ΔΔG| and RSA**
The figure shows the correlation between |ΔΔG| (y-axis) and Relative Solvent Accessibility (RSA, x-axis). The ancestral RSA is the RSA in the ancestral human Alphafold structure, estimated in the Thermonet framework. The blue line shows the linear regression fit. The negative correlation is significant ($P<10^{-16}$).

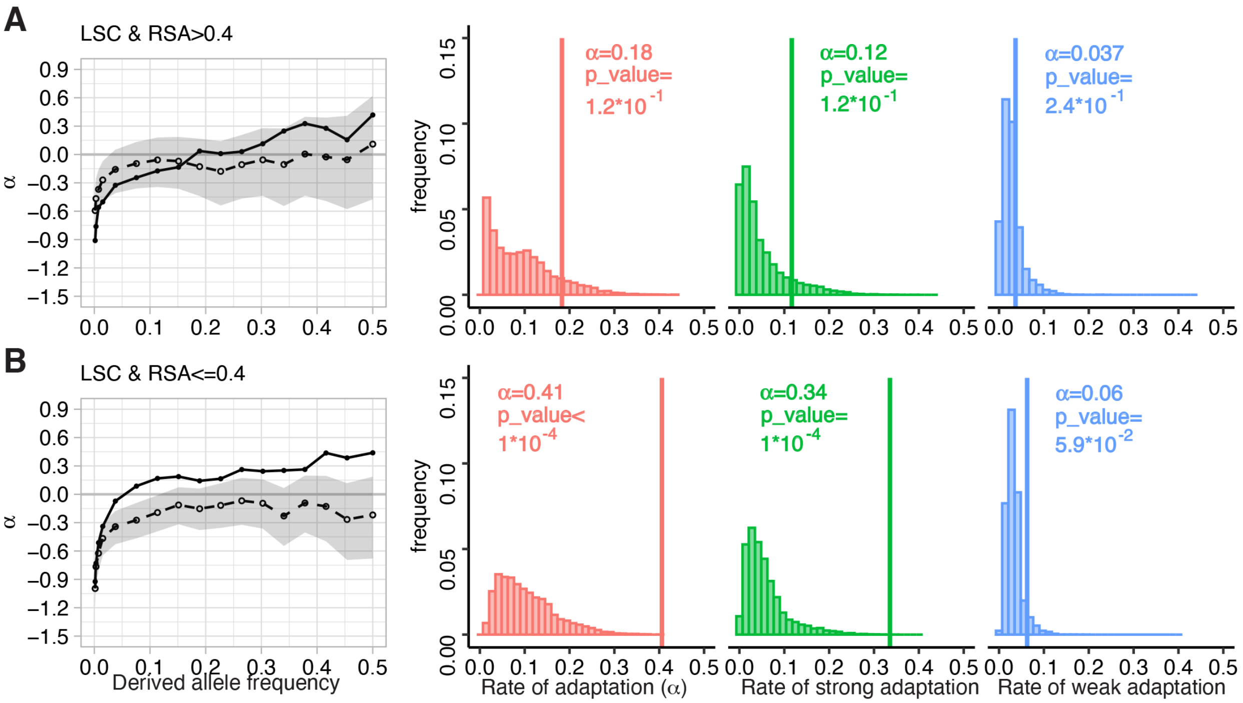


**Figure S2. Adaptation with large stability changes as a function of RSA**
A and B legend same as Figure 3C,D. The difference between this figure and Figure 3C,D is that in Figure 3C,D, the RSA value (0.23) to split between high and low RSA is the median among LSCs. However the value to split between high and low RSA for SSCs (Figure 3E,F) is 0.4, the median RSA for SSCs. Figure S2 uses the same RSA=0.4 split value between A and B, and is therefore more directly comparable to Figure 3E,F than Figure 3C,D.

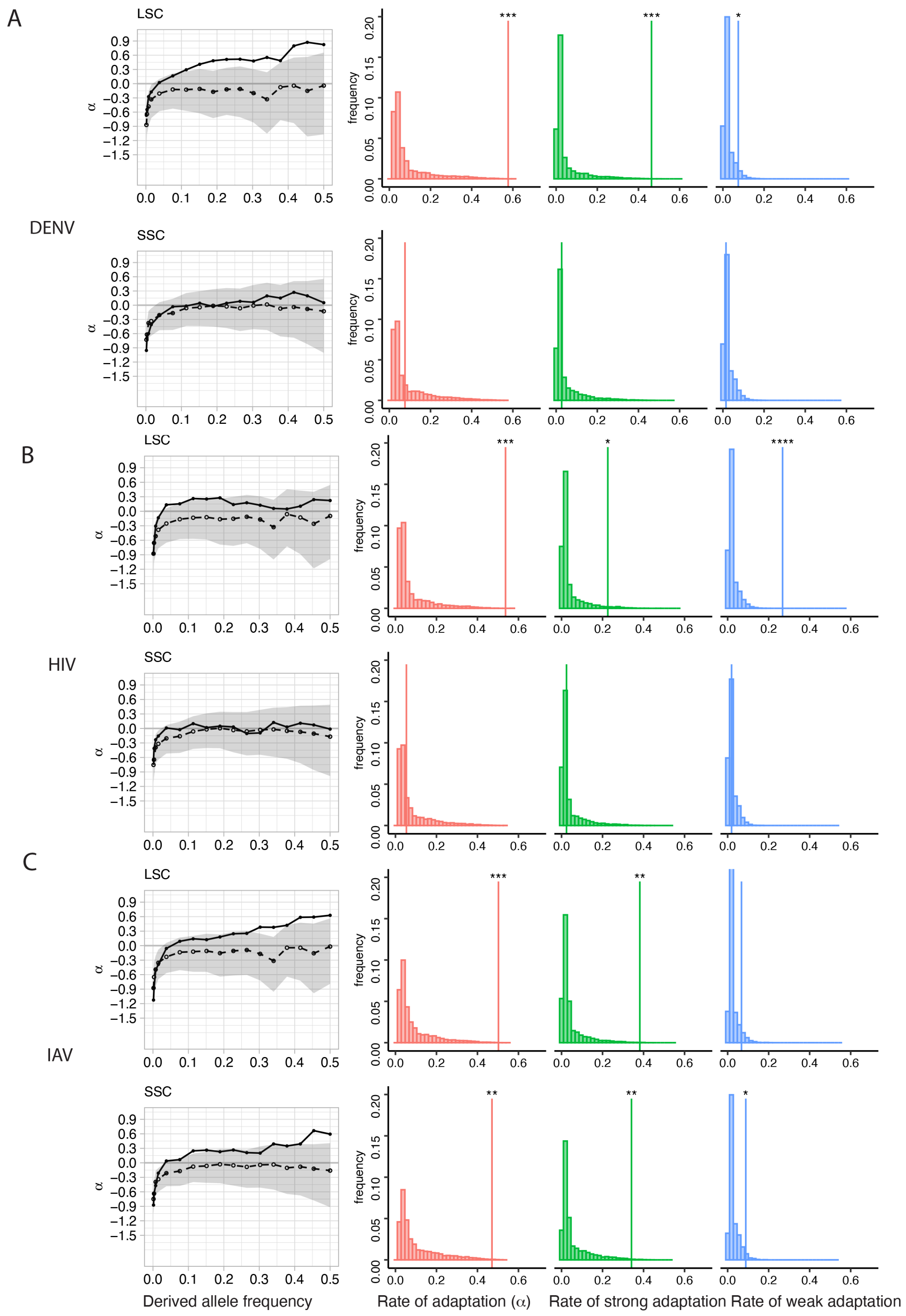


**Figure S3. Adaptation with LSCs or SSCs in DENV, HIV, or IAV VIPs**

Same legend as Figure 4.
